# The cortex encodes speech timing as departure from expectation across multiple timescales

**DOI:** 10.64898/2026.08.19.745735

**Authors:** Nicola Molinaro, Jose Pérez-Navarro

## Abstract

Every syllable and every phoneme in natural speech has a different duration. This variability is conventionally treated as jitter, noise the brain must overcome to recover an underlying regularity. Here we show that the cortex encodes it as information. We recorded magnetoencephalography from 25 native listeners of Spanish during twenty minutes of spontaneous narrative speech in Spanish, a syllable-timed language in which durational variability is low, making it a conservative test case. We modelled cortical activity with three timing regressors defined at successively finer grains: deviation from the expected syllabic beat, from the expected phoneme onset, and from the expected duration of the phoneme just completed. Fitted jointly, and against acoustic and linguistic-surprisal baselines, each regressor explained unique activity in bilateral superior temporal cortex, with distinct response latencies spanning 70 to 250 ms. The duration regressor explained cortical activity beyond raw phoneme duration, whereas raw duration explained no additional variance once it was included. At the onset grain, by contrast, predicted and physical timing contributed jointly. The onset departure was encoded with its sign, distinguishing events that arrived early from those that arrived late. In a separate sample, removing this durational variability was costly: comprehension in noise degraded under imposed isochrony relative to natural timing. Durational variability is therefore not noise the cortex discards but information it encodes, across a hierarchy of timescales — and at the finest grain, what is encoded is not how long a speech event lasted but how far its duration departed from expectation.

---

A listener who is to anticipate what is about to be said must exploit the temporal structure of the unfolding signal. A substantial body of evidence indicates that the cortex does so by entraining its endogenous activity to the slow, quasi-periodic envelope of syllabic rhythm^1–3^. Speech rhythms, in this broad sense, constitute one dimension along which the brain organises auditory cortical processing, and the case for their role is robust. Yet even at the syllabic level, speech is only quasi-periodic: successive syllables vary substantially in duration, and the variability is not noise around a stable rate but lawful structure that reflects phonological category, stress, and prosodic position. A neural mechanism that tracks regularity therefore captures only the global temporal scaffold of the signal it must process. The central computational problem is not how the brain locks onto regularity, but how it derives information from variability.

If rhythm-tracking is to suffice as an account of speech-timing processing, it should suffice most readily in those languages whose surface timing is closer to periodicity. Spanish, canonically classified as syllable-timed (a rhythmic class whose amplitude-modulation spectra are distinguishable from those of stress-timed languages^4^), is the natural test case. Yet even in Spanish the temporal structure of natural speech departs substantially from periodicity. Successive inter-onset intervals between syllables and between phonemes vary widely from one event to the next, and this variability is unevenly distributed across timescales. The variability is too pronounced for a single-frequency tracker to lock onto, and too systematic to be dismissed as motor noise around an underlying periodicity.

The departure from periodicity is, on its own, only a problem for accounts that *expect* periodicity; it does not yet establish what the brain is doing instead. Two further observations make the point sharper. First, when speech is artificially flattened to perfect isochrony, the condition that should be optimal for an oscillator tuned to syllabic rate, listeners’ comprehension worsens rather than improves, as observed in speech-in-noise experiments^5,6^. Across natural speech, comprehension is better when periodicity is lower^7^. Second, phoneme durations are not free to vary. They are structured by phonological category, lexical stress and prosodic position^8–10^, and further modulated by informativeness, speakers lengthening unpredictable material and compressing redundant material^9,10^. What an oscillator would treat as noise is, on inspection, lawful. Taken together, these observations imply that timing variability is not an obstacle the comprehension system overcomes but a signal it exploits. That listeners exploit lawful variability is not itself a new proposal: contextually conditioned variation in the speech wave has long been argued to be used in recovering the intended segments rather than discarded as distortion^11^. But that argument, like the production-side evidence above, concerns which sounds are spoken. Recent work has begun to extend it to when they occur: the onsets of phonemes, syllables and words in natural speech are predictable from the timing of preceding events alone, and the resulting predictions explain intracranial responses beyond acoustic and linguistic content^12^. What form that encoded quantity takes, however, remains open. There, timing predictability was best captured by the untransformed probability of an upcoming onset rather than by its surprisal — a transformation that, being a function of probability alone, is blind to whether an event arrived early or late. A mechanism that exploits timing variability rather than suppressing it must be *adaptive*, in that it adjusts moment-to-moment to a non-stationary signal; *local*, in that it operates at the timescale of individual phonological events rather than over global statistics; and *variability-sensitive*, in that it treats deviations from expectation as carriers of information rather than as error to be suppressed.

The three properties identified above do not jointly describe a generic mechanism; they jointly describe a predictive one. Variability cannot be tracked as such: a deviation is defined only with respect to an expectation, and a mechanism that is sensitive to deviations is, by construction, a mechanism that generates predictions. To be adaptive and local is to revise those predictions continuously and at the timescale of events; to be variability-sensitive is to compute the residual between predicted and observed. These are not three properties that *happen* to characterise a predictive system among others. They are constitutive of one. On this view, the cortical response to natural speech is shaped not by the timing of events as such but by the deviation of each event’s timing from a moment-to-moment expectation generated from recent context. Stated this way, the account requires a corresponding change in what we measure. Speech neuroscience has asked how the cortex extracts regularity from a variable signal, treating the residual variability as what the analysis must factor out. The inversion is exact: the residual is the quantity encoded, event by event, and it is measurable as such.

The sharpest test of this proposal concerns the cortical representation of phoneme duration — a quantity that onset-prediction accounts do not address, since it is defined only once an event has ended. If the brain represents the physical timing of events, then the duration of each phoneme in milliseconds, a property of the acoustic event itself and independent of any prediction, should drive a cortical response. If the brain represents only deviation from predicted timing, then raw duration should explain no cortical variance once its predictive transformation, computed under a class-conditioned model of expected duration, is accounted for. The two accounts make opposite empirical predictions, and their contrast can be evaluated within a single statistical model.

We test these proposals in Spanish, for the reason given above. We recorded magnetoencephalography from twenty-five native Spanish listeners during twenty minutes of continuous spontaneous speech, force-aligned at the phoneme level. From these we derive three families of timing-prediction regressors, corresponding to three candidate levels of adaptive temporal prediction: syllabic rhythm, sub-syllabic phoneme onset, and phoneme duration (**Fig. 1**). We fit them together with acoustic and linguistic predictors^13,14^ in a single temporal response function model in the superior temporal cortex. The results, presented below, indicate that adaptive temporal prediction at multiple timescales operates in parallel with linguistic prediction, and is statistically independent of it, and that the cortical response to phoneme duration is exhausted by its predictive transformation.

**Figure 1.**
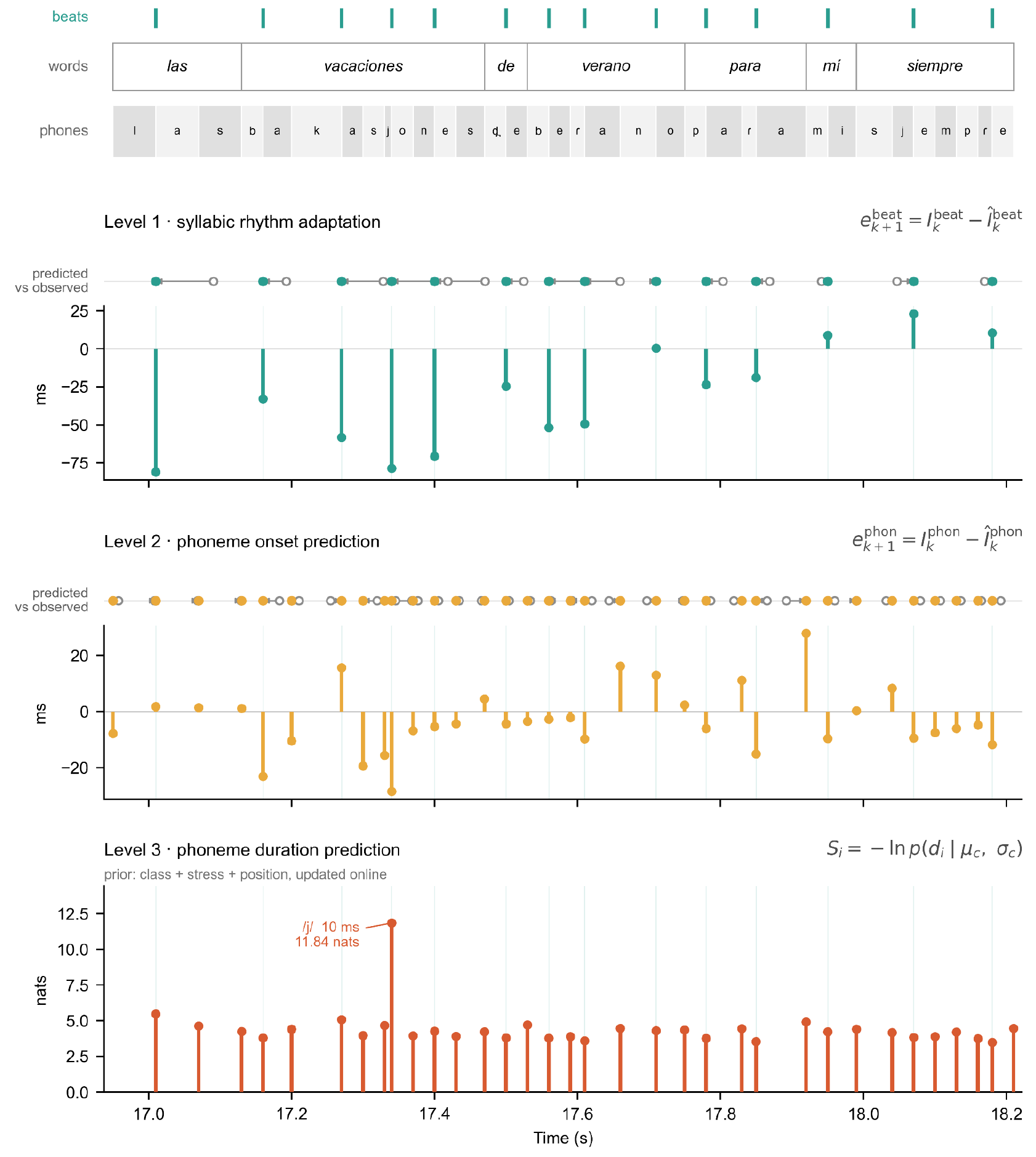
Construction of the three timing regressors. A 1.28-s excerpt from one narrative (“*…las vacaciones de verano para mí siempre…*”), showing the regressor values actually entered into the temporal response function models. Top, word and phoneme boundaries from forced alignment; teal ticks mark vowel onsets, taken as the syllabic beat. Levels 1–3, each level applies the same predict-and-compare operation at a different linguistic grain. For Levels 1 and 2, the strip above each panel shows the predicted event time (open circle), formed from an exponential moving average of the inter-onset intervals, against the observed event time (filled circle); the arrow is the deviation between them, which is read out as the regressor value below. Negative values indicate an event arriving earlier than predicted, positive later. Level 1 is defined at vowel onsets and its value is held constant across intervening consonants, so the regressor changes only at syllabic nuclei; Level 2 assigns each phoneme its own onset deviation over the full segmental sequence. Both are expressed in milliseconds and entered as impulses at phoneme onsets. Level 3 is entered at phoneme offsets and expressed in nats, as the surprisal of the realised duration under a class-conditioned expectation updated online after every phoneme; because it compares a duration rather than an event time, it has no predicted-versus-observed strip. Annotated is the /j/ of *vacaciones*, realised in 10 ms against a median glide duration of 40 ms in this narrative, yielding 11.84 nats — the largest duration surprisal in the excerpt.

## Results

### Natural speech timing aids comprehension in noise

If listeners exploit the temporal variability of speech rather than merely tolerating it, then removing that variability should cost them. We tested this behaviourally in a group of native Spanish listeners (N=27). Listeners heard sentences embedded in white noise (−1 dB SNR) under three timing conditions: natural timing preserved, timing flattened towards isochrony, and timing exaggerated into anisochrony (irregularity). Keyword recall differed across conditions (one-way repeated-measures ANOVA, F(2, 52) = 940.86, p < .001, η²ɢ = 0.67; Mauchly’s test indicated no violation of sphericity, W = 0.86, χ²(2) = 3.75, p = .154). Recall was highest for natural timing (80.6 ± 7.8%), lower under isochrony (62.1 ± 10.1%), and lowest under exaggerated irregularity (48.8 ± 9.8%). All pairwise contrasts were significant (Bonferroni-corrected paired t-tests, all df = 26): natural exceeded isochronous by 18.5 percentage points (95% CI [17.2, 19.9], t = 27.97, p = 1.9 × 10⁻²⁰, d_z = 5.38), and anisochronous by 31.8 points (95% CI [30.0, 33.6], t = 36.85, p = 1.7 × 10⁻²³, d_z = 7.09); and isochronous exceeded anisochronous by 13.3 points (95% CI [11.9, 14.7], t = 19.93, p = 8.5 × 10⁻¹⁷, d_z = 3.84; **Fig. 2**). Comprehension therefore tracked how closely timing matched the statistics of natural speech, not how regular it was. This ordering replicates findings in French, English and Australian English^5,6^, and here extends to Spanish. The benefit of natural variability is not tied to a single language.

**Figure 2.**
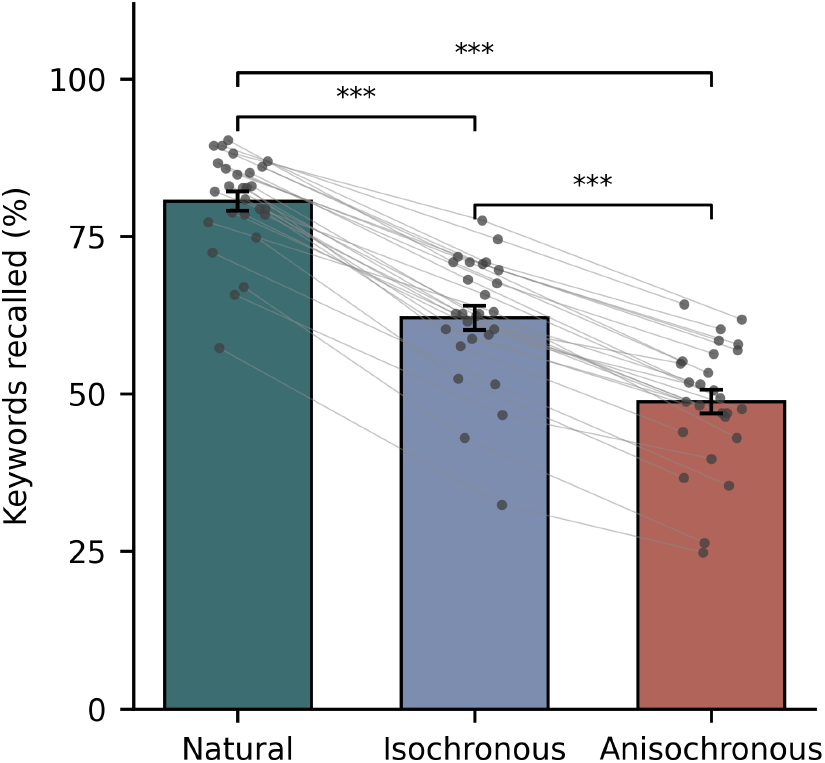
Keyword-recall accuracy for sentences presented in white noise (−1 dB SNR) under three timing conditions — natural, isochronous (timing flattened to regularity), and anisochronous (timing exaggerated into irregularity) — for N = 27 listeners. Recall was highest for natural timing (80.6 ± 7.8%, mean ± SD), lower under isochrony (62.1 ± 10.1%), and lowest under anisochrony (48.8 ± 9.8%); all pairwise differences were significant (repeated-measures ANOVA, F(2, 52) = 940.86, p < .001, η²ɢ = 0.67; Bonferroni-corrected paired t-tests, all df = 26, all p < 10⁻¹⁶). Trials on which participants produced no response were scored as zero keywords recalled. Bars show group means, error bars ±1 SEM, dots individual participants.

### Three predictive-timing regressors derived from continuous speech

To ask whether the auditory cortex encodes speech timing predictively, we derived three families of regressors. All three express the same logic: a prediction of when or how long a speech event should be, formed from local context, against which the observed event is compared. They differ in the linguistic grain at which that prediction is cast. The *first* operates at the syllabic scale: from the recent rhythm of vowel onsets it anticipates when the next syllabic beat should fall, and encodes the discrepancy when it falls early or late. This level is indifferent to which syllable arrives; it registers only temporal regularity and its violation. The *second* applies the same predict-and-compare operation at the finer, non-rhythmic grain of individual phonemes, anticipating the onset of the next segment from the local pace of articulation. It carries minimal linguistic commitment: it tracks segmental timing without regard to segmental identity. The third is explicitly linguistic. For each phoneme it holds a category-conditioned expectation of duration, how long a sound of this class in this prosodic context should last, and reads out the surprisal of the realised duration under that expectation. Across the three, the amount of linguistic knowledge the estimator presupposes increases — minimal, requiring only the location of syllabic nuclei, then the segmentation of the signal into phonemes, then their phonological and prosodic classification — while the organising principle remains constant: expectation followed by comparison with observed timing.

We recorded MEG from 25 native Spanish listeners as they attended to continuous spontaneous speech. Each of five narratives was a speaker talking for roughly seven minutes about a specific topic; every participant heard three, drawn pseudo-randomly and counterbalanced across participants, for some twenty minutes of listening in total. After each narrative, participants answered comprehension questions; accuracy was high (mean 93.9%, SD 7.6), confirming that they attended to the speech throughout. Spontaneous speech maximised naturalness. It also raises an immediate question about the timing statistics of the signal itself. Spanish is conventionally classed as a syllable-timed language, the very type expected to approximate temporal regularity most closely; if any natural speech were periodic enough to sustain an account based on entrainment to a fixed beat, it should be this. It is not. Across the five narratives, inter-onset intervals were highly variable at both grains we consider: the coefficient of variation^15^ reached ≈0.76 for syllabic onsets and ≈1.10 for phonemes, placing both well outside the near-isochronous regime a fixed oscillator would require. The normalised pairwise variability index, computed here over inter-onset intervals rather than the vocalic intervals for which it was originally defined^16,17^, was of comparable magnitude at the two scales (≈51 at both grains). The irregularity is not confined to a single timescale (**Table 1**). Natural speech of this kind is poorly described as a carrier with a stable period. Its timing is structured but not periodic: the regime in which prediction, rather than entrainment, becomes necessary.

**Table 1.** Temporal regularity of the five narrative segments. Coefficient of variation (CV) and normalised pairwise variability index (nPVI) of inter-onset intervals, computed separately at the phoneme grain and the syllabic grain. Silent intervals were removed and each segment was split at pauses ≥150 ms, with intervals computed within the resulting runs. Syllabic intervals were taken between successive vowel nuclei; adjacent vowels were treated as a single nucleus where one member was a high vowel (/i/, /u/) and as separate nuclei otherwise. Counts are of intervals rather than events.

| Segment | Phonemes (n) | Phoneme CV | Phoneme nPVI | Syllables (n) | Syllable CV | Syllable nPVI |
| --- | --- | --- | --- | --- | --- | --- |
| 1 | 5067 | 1.24 | 50.1 | 2049 | 0.83 | 48.9 |
| 2 | 4496 | 1.04 | 51.3 | 1809 | 0.76 | 52.7 |
| 3 | 4357 | 1.12 | 52.9 | 1782 | 0.77 | 56.0 |
| 4 | 4305 | 1.11 | 51.7 | 1713 | 0.77 | 53.1 |
| 5 | 4970 | 1.01 | 49.6 | 2020 | 0.70 | 50.2 |
| **Mean ± SD** | — | **1.10 ± 0.09** | **51.1 ± 1.3** | — | **0.76 ± 0.05** | **52.2 ± 2.7** |

Before relating these regressors to cortical activity, we confirmed they are not proxies for linguistic-content predictions. Across all five narratives (24,138 phonemes; **Fig. 3**), the three timing predictors were near-independent of sublexical, word, and sentence-level surprisal: every timing-by-surprisal correlation fell at or below r = 0.17, and phoneme-duration surprisal (Level 3) was effectively orthogonal to all three (|r| ≤ 0.03). Within the timing family the levels were only modestly related (largest r = 0.22, between phoneme-onset and phoneme-duration prediction), as were the linguistic predictors among themselves in the expected nested pattern. Timing and linguistic predictions thus occupy largely separate regressor subspaces. From these regressors, together with acoustic and linguistic baselines, we estimated source-localised temporal response functions in the superior temporal cortex.

**Figure 3.**
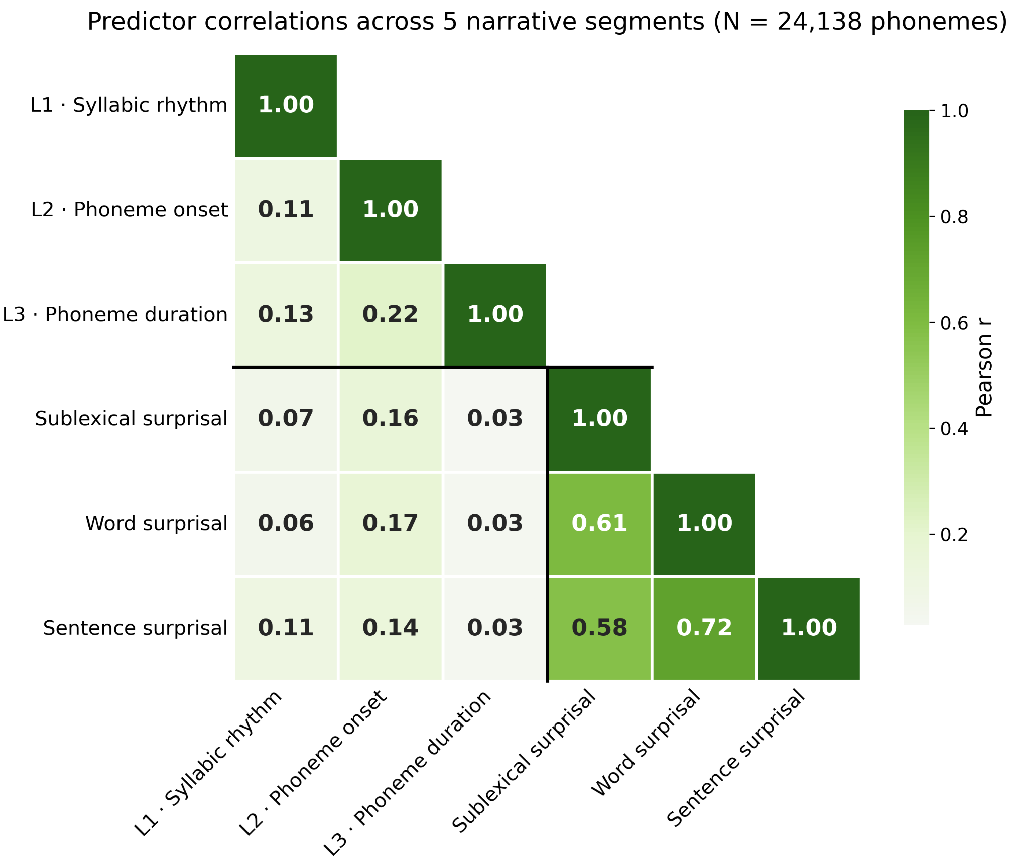
Timing and linguistic predictors occupy separate subspaces. Pearson correlations among the three predictive-timing regressors (Level 1, syllabic rhythm; Level 2, phoneme onset; Level 3, phoneme duration) and the three linguistic-surprisal regressors (sublexical, word, sentence), computed across all five narrative segments (N = 24,138 phonemes). The three timing predictors are near-independent of the linguistic predictors (all |r| ≤ 0.17), and phoneme-duration surprisal (Level 3) is effectively orthogonal to them (|r| ≤ 0.03); within the timing family the levels are only modestly related (largest r = 0.22, Levels 2–3). The black divider separates the timing block (upper left) from the linguistic block (lower right).

### The cortex encodes phoneme duration as departure from expectation, not as physical length

We fit temporal response functions relating each set of regressors to source-localised MEG within bilateral superior temporal gyrus^13^, and compared nested models by the improvement in predictive accuracy each regressor afforded (threshold-free cluster enhancement (TFCE), described in *Statistical analysis of model comparisons*, 10,000 permutations).

We first asked whether the cortical response to each phoneme reflects its physical duration or its departure from a category-conditioned expectation. Raw duration and duration surprisal can be entered at the identical event time, the phoneme offset, and so can be compared with timing held constant. The TRF model contained acoustic (gammatone spectrogram and gammatone onsets), a phoneme-onset impulse and linguistic-surprisal regressors, the Level-1 and Level-2 deviation predictors, raw phoneme duration, and duration surprisal. Raw phoneme duration, added at the offset, explained no additional cortical activity beyond the rest of predictors (t_max = 1.44, p = .815, df = 24). The reverse contrast was significant: duration surprisal, entered at the same offset, improved model fit over a baseline already containing raw duration alongside the same set of predictors (t_max = 3.79, p = .024, df = 24). Because event timing is matched across the two regressors, the only difference between them is the predictive transformation. The cortical response to phoneme duration is therefore carried by its departure from a category-conditioned expectation, not by physical duration itself.

### Adaptive temporal prediction operates at three timescales

Having shown that the cortical response to duration is predictive rather than physical, we asked whether the same holds at the two coarser levels, and whether all three contribute independently. Each timing predictive regressor explained cortical activity beyond the acoustic and linguistic-surprisal baseline and the other two timing regressors (bilateral superior temporal gyrus, TFCE, 10,000 permutations, one-tailed). The syllabic-beat prediction (Level 1: t_max = 3.87, p = .002), the phoneme-onset prediction (Level 2: t_max = 6.76, p < .001), and the phoneme-duration surprisal (Level 3: t_max = 3.60, p = .015) respectively improved model fit over a baseline already containing gammatone spectrogram, gammatone onsets, phoneme onsets, and sublexical, word, and sentence-level phoneme surprisal (all df = 24). Each test evaluates a distinct, pre-specified level of the architecture and is reported individually; all three remain significant under Bonferroni correction across the three comparisons (corrected p = .006, <.003 and .045 for Levels 1–3). Predictive timing is therefore a separable dimension of the cortical response, operating in parallel with linguistic prediction at three distinct timescales.

To quantify effect magnitude and cross-participant consistency, which the mass-univariate test does not provide, we extracted each participant’s Δdet averaged over the bilateral superior temporal mask. Note that averaging over the full mask dilutes spatially restricted effects, so these values index magnitude and consistency rather than providing an independent test. Level 1 improved prediction by Δdet = 4.67 × 10⁻⁵ (95% CI [1.79, 7.54] × 10⁻⁵; t(24) = 3.35, p = .003, d_z = 0.67; 18/25 participants positive). Level 2 improved it by 7.40 × 10⁻⁵ (95% CI [4.10, 10.70] × 10⁻⁵; t(24) = 4.62, p < .001, d_z = 0.92; 23/25). The Level 3 difference was smaller and more variable (Δdet = 6.27 × 10⁻⁶, 95% CI [−0.05, 1.30] × 10⁻⁵; t(4) = 1.92, p = .067, d_z = 0.38; 14/25) and did not reach significance when averaged over the full mask. The Level 3 contribution was reliable in the mass-univariate test (t_max = 3.60, p = .015) but spatially restricted.

The three levels differed in response latency and shape (**Fig. 4**). The Level 1 response rose slowly to a broad maximum at 200 ms in the left and 190 ms in the right hemisphere. This latency is in the range of the syllabic interval itself, consistent with a response that accumulates across a syllabic cycle rather than reacting to a discrete acoustic edge. The Level 2 response was biphasic, with a negative deflection reaching its trough at 60–70 ms after nominal phoneme onset, followed by positive deflections at approximately 110, 230–260 and 290–340 ms. Some pre-onset activity is expected for phoneme-locked responses given coarticulation and the quasi-periodic structure of phoneme onsets. The Level 3 regressor is time-locked to phoneme offset rather than onset. Its sharp peak at 70 ms in both hemispheres therefore indexes a response arising shortly after a phoneme has ended, when its realised duration first becomes available to the listener. Peak latencies were near-identical across hemispheres for all three levels, but response amplitudes were not. The Level 1 response was larger over the left hemisphere throughout its rise and peak (160–190 ms, p = .025; 210–260 ms, p = .009; 300–320 ms, p = .025; peak t = 3.00–3.38), with a later cluster in the opposite direction at 520–550 ms (t = −4.07, p = .005). The Level 2 response was larger over the left hemisphere on the trailing edge of its third positive deflection (340–370 ms, t = 4.10, p = .005), and Level 3 showed no hemispheric difference. These amplitude asymmetries did not translate into asymmetries in predictive contribution, which did not differ statistically between hemispheres at any of the three levels (Level 1, t_max = 2.71, p = .168; Level 2, t_max = 3.88, p = .056; Level 3, t_max = −0.94, p = .573).

**Figure 4.**
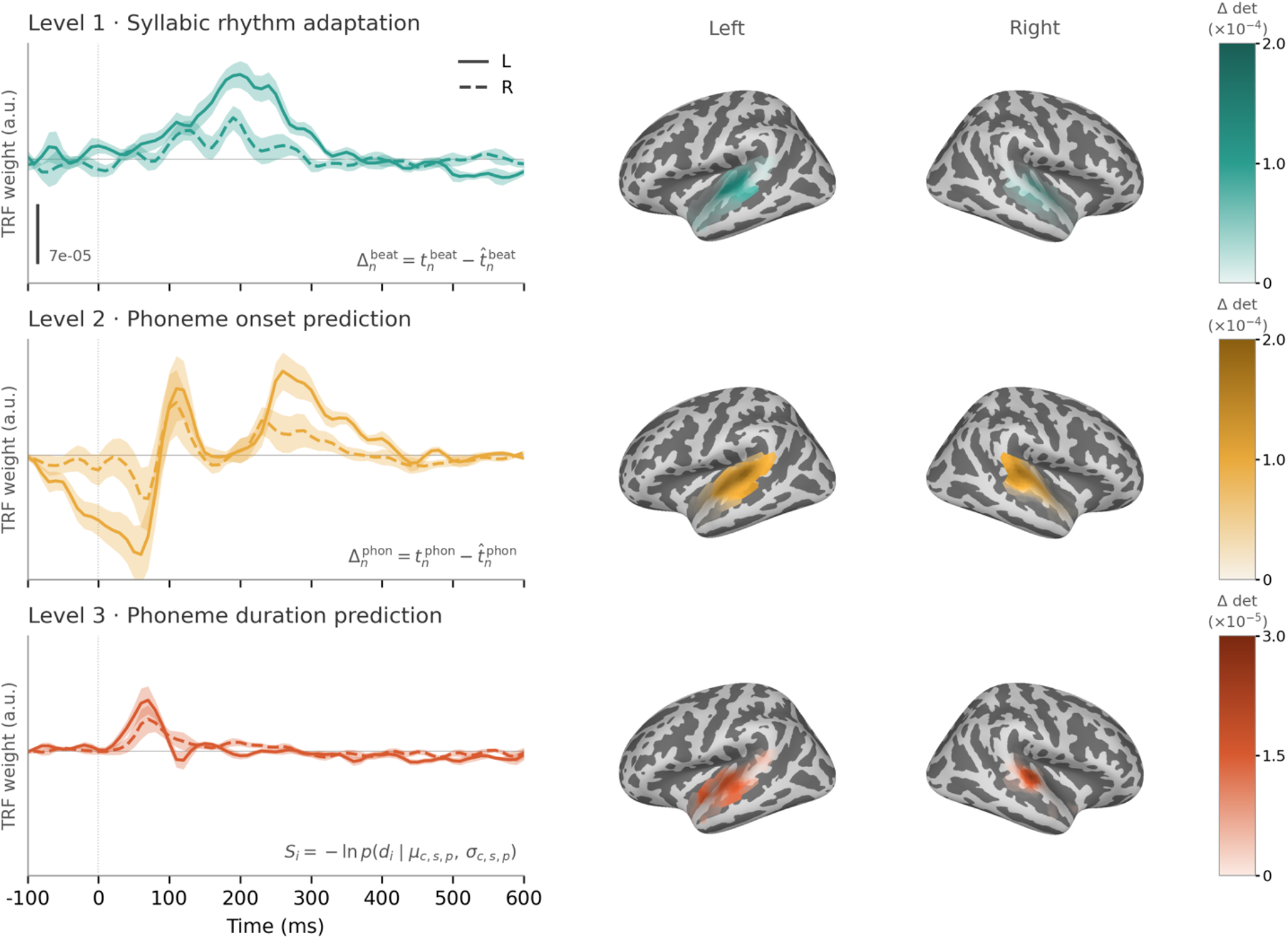
Cortical responses to the three levels of temporal prediction. *Left*, temporal response functions (TRFs) for each timing regressor, averaged over sources in the bilateral superior temporal mask and plotted separately for the left (solid) and right (dashed) hemisphere; shading denotes within-subject standard error across the 25 participants. Weights are in arbitrary units, as responses and predictors were standardised before boosting; the vertical bar in the top panel calibrates all three panels, which share a common ordinate. Level 1 and Level 2 regressors were both represented at phoneme onsets. Level 1 carried forward the most recent vowel-onset departure and changed only at vowel onsets, whereas Level 2 carried the departure calculated from the complete phoneme-onset sequence. Level 3 was represented at phoneme offsets. Insets give the corresponding quantity. *Right*, increase in predictive power (Δdet) attributable to each regressor, computed as the difference between the full model and the same model with that regressor omitted, thresholded at p ≤ 0.05 (TFCE, 10,000 permutations) and rendered on the inflated *fsaverage* surface. Colour scales differ by level; note the exponent on each bar.

Comparing a model carrying the three timing regressors with one carrying the full sublexical/word/sentence surprisal hierarchy, we found no reliable difference (t_max = 2.38, p = .601). We do not read this as evidence of equivalence. What the data do establish is that each family contributes uniquely in the full model: encoding when speech events occur is not redundant with encoding what is said.

### The onset departure is signed, and does not exhaust the physical interval

The Level-2 regressor is signed: negative when a phoneme arrives earlier than expected, positive when later. We asked whether the cortex is sensitive to that sign or only to the size of the departure. It is sensitive to the sign. Adding the direction of the departure to a model already containing its magnitude improved model fit (t_max = 3.23, p = .020), and a direct comparison of the signed and unsigned forms favoured the signed one (t_max = 3.62, p = .018). Splitting the departure into its two directions, the late-arrival regressor explained cortical activity beyond a model already containing the early-arrival regressor (t_max = 5.52, p < .001): the two directions are not interchangeable. What the cortex encodes at this grain is not only how far an event departs from its expected time, but in which direction. The two directions were carried by responses at different latencies. The early-arrival response peaked at 60 ms after nominal phoneme onset and the late-arrival response at 230–250 ms, and the two curves differed reliably over the left hemisphere from −50 to 80 ms and again from 200 to 400 ms (two-tailed, TFCE, 10,000 permutations). The same pattern was present over the right hemisphere at lower amplitude, where it did not reach significance. The biphasic Level 2 response described earlier (**Fig. 4**) is therefore not a single response with two phases but the superposition of two: an early deflection driven by phonemes arriving sooner than expected, and a later one driven by phonemes arriving late (**Fig. 5**).

**Figure 5.**
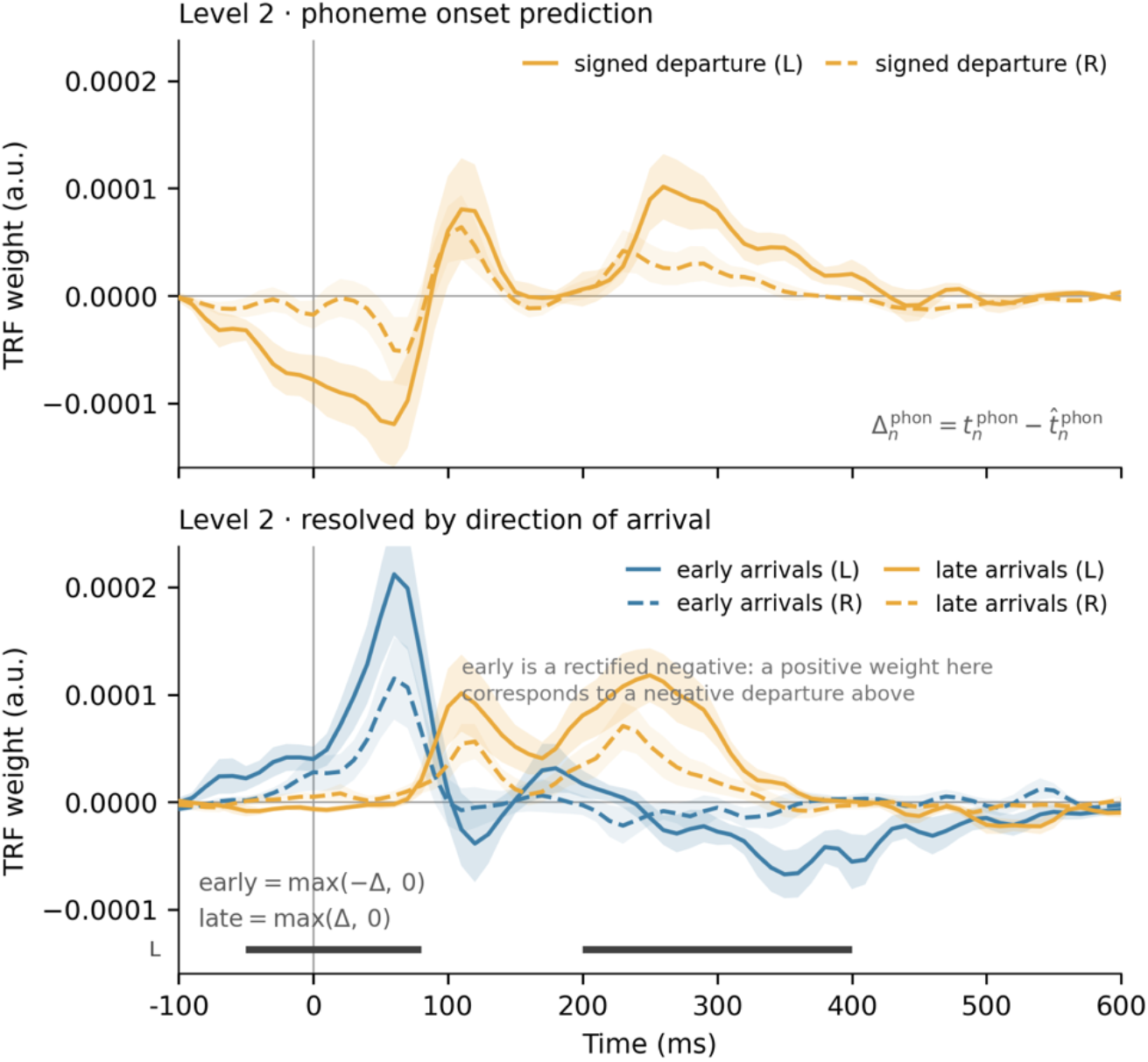
The Level 2 response resolves into two responses with different latencies. Temporal response functions for the phoneme-onset predictors, averaged over sources in the bilateral superior temporal mask and plotted separately for the left (solid) and right (dashed) hemisphere; shading denotes within-subject standard error across the 25 participants. Weights are in arbitrary units, as responses and predictors were standardised before boosting, and the two panels share an ordinate. *Top*, the signed Level 2 departure, the same regressor plotted in the Level 2 row of Figure 4. *Bottom*, the departure resolved into its two directions: an early-arrival regressor carrying the magnitude of the departure when a phoneme arrived sooner than expected and zero otherwise, and a late-arrival regressor defined symmetrically for later-than-expected arrivals. Each event activates exactly one of the two, so the pair is disjoint by construction and the difference in latency between them cannot arise from variance shared between the regressors. The early-arrival response peaks at 60 ms and the late-arrival response at 230–250 ms; the horizontal bars mark the windows in which the two differ over the left hemisphere (two-tailed, TFCE, 10,000 permutations, p < .05). Because the early-arrival regressor is a rectified negative, a positive weight in the lower panel corresponds to a negative weight on the signed departure above. The two panels come from separate fits of the same data, each regularised independently, so the two directional responses are not expected to sum exactly to the response above them.

We then asked whether the departure exhausts the physical interval, as it does at the duration level. It does not. The Level-2 departure explained cortical activity beyond the raw inter-onset interval (t_max = 5.70, p < .001), but the raw interval likewise explained activity beyond the departure (t_max = 5.58, p < .001), and a non-nested comparison of the two models showed no reliable difference (t_max = 2.81, p = .253). The two regressors are correlated at r = 0.68, so this is not a case of two independent quantities each finding its own variance: each adds over the other despite substantial overlap. At the onset grain, predicted and physical timing contribute jointly; neither supersedes the other. The asymmetry reported above for duration — where raw duration added nothing once its surprisal was included, while the reverse contrast was significant — is therefore specific to Level 3.

## Discussion

### Multi-level, adaptive prediction of event timing

Our results indicate that the auditory cortex encodes the timing of speech events predictively. The three timing regressors captured cortical activity that neither the acoustic signal nor linguistic surprisal could explain. Each formalises a prediction of when or how long an event should be: the syllabic beat, the phoneme onset, and the category-conditioned duration of each phoneme. These operate in parallel with, and separably from, the prediction of linguistic content.

Timing prediction is not a by-product of anticipating upcoming words or phonemes, but a distinct axis of the cortical response. We do not claim to have shown that variability-sensitivity logically entails prediction. That inference is a theoretical premise, and what our data test are its consequences: most directly, that the cortical response to phoneme timing is governed by deviation from expectation.

The three levels differ in grain and in what they commit to. The syllabic level is the coarsest: it tracks the pace of speech, an evolving expectation of when the next syllabic beat should fall, indifferent to which syllable arrives. The phoneme-onset level applies the same logic at a finer, non-rhythmic grain, anticipating when the next segment is due. The duration level is the finest and the only one that is linguistically specific, holding a category-conditioned expectation of how long each phoneme should last. The three address complementary questions — the pace, the *when*, and the *how long* — yet share a common principle: a locally formed expectation against which observed timing is evaluated. Each contributed independently to the cortical response; timing, in natural speech, is predicted at all three grains in parallel.

The departure our regressors encode is signed, and the cortex is sensitive to that sign. Adding the direction of a phoneme’s onset departure improved model fit over its magnitude alone, and separating early from late arrivals showed the two are not interchangeable. They were carried by responses at different latencies, and the biphasic shape of the Level 2 response resolves into the superposition of the two. This matters for what kind of quantity is being represented. A departure that carries only magnitude is a measure of how badly an expectation failed; a signed departure also says in which direction the world diverged from the model, and so remains informative about the event itself, not only about the quality of the prediction. Recent work reaching the same problem from onset timing found that the untransformed probability of an upcoming onset explained cortical activity better than its surprisal^12^. Surprisal is a function of probability alone, and is therefore blind to direction: an event arriving early and one arriving late are assigned the same value. Our result gives a reason why a surprisal transform might underperform on this quantity. It is not that timing departures go unencoded, but that collapsing them onto a magnitude discards information the cortex retains.

A second result at this level was not predicted. The raw inter-onset interval explained cortical activity beyond the departure, just as the departure explained activity beyond the interval, and the two models were statistically indistinguishable when compared directly. At Level 3 the corresponding contrast was one-sided: raw duration added nothing once its surprisal was included. The asymmetry is worth taking seriously rather than explaining away, and we offer the following reading post hoc. The departure is computed from the interval: it is the difference between the observed interval and an expectation formed from preceding ones. A system that encodes the departure must therefore have the interval available, and finding both encoded is what a single computational pipeline would produce if its input remains accessible alongside its output. Level 3 differs in that its input is a completed duration, a quantity with no evident further use once its surprisal has been read. On this reading the two levels are not in conflict: what is exhausted by the predictive transformation is the measurement that the transformation consumes, and the onset interval is not consumed. We note that this account was formed after the result and requires an independent test.

Converging evidence for that separation comes from outside speech: rhythm-based and interval-based temporal prediction can be doubly dissociated, being differentially impaired in Parkinson’s disease and in cerebellar degeneration^18^. The interval-based prediction studied there is cued and single-shot rather than continuously updated. Even so, the dissociation supports treating rhythmic and non-rhythmic timing prediction as distinct mechanisms rather than one mechanism read out at two grains.

A limitation should be acknowledged at the third level. The Level 3 contribution, while significant, was the weakest of the three (p = .015). The duration-versus-surprisal contrast provides convergent evidence from a different test design: duration surprisal explained cortical variance that raw duration could not (p = .024), confirming that the cortex encodes phoneme duration predictively. Replication in an independent sample and language will be important to establish the robustness of this level.

The sharpest form of this claim concerns duration. Raw phoneme duration, entered in milliseconds, explained no cortical variance once its predictive transformation was included: the surprisal of that duration given the phoneme’s category and context. Reversely, phoneme duration surprisal did explain cortical activity beyond raw phoneme duration. What the cortex encodes is therefore not how long a sound lasts, but how far its length departs from what was expected. We speculate that this reflects a more general principle. Time may not be a property the stimulus carries but, as has been argued^19^, an abstracted relational measure of change: here, the relation between when an event occurs and when it was predicted to occur.

The Level 1 response was larger over the left hemisphere through its rise and peak, although its contribution to predictive accuracy was bilateral. This is worth noting because accounts of asymmetric temporal sampling associate the syllabic timescale with a right-hemisphere bias^20,21^, which is what an envelope-tracking mechanism would predict. Our Level 1 regressor is not an envelope tracker: it carries the departure of each syllabic interval from a locally formed expectation, and its left-weighted response is more consistent with a quantity feeding linguistic analysis than with acoustic sampling. We advance this cautiously — a between-hemisphere amplitude difference in fixed-orientation source estimates is not a clean index of lateralised computation, and the predictive-accuracy measure showed no such asymmetry.

### Temporal variability as information, not noise

The temporal structure of natural speech is irregular: in our materials, inter-onset intervals varied widely at both the syllabic and the phonemic grain, far from the isochrony that a fixed oscillator would require. On a view in which the brain’s task is to recover a periodic carrier, this variability is a nuisance: jitter to be smoothed away before the signal can be read. Our findings point the other way. Listeners understood speech best when its natural timing was preserved and worst when that timing was regularised or exaggerated. Cortical activity was structured by the moment-to-moment departure of each event from what was predicted. The variability is not noise around a signal; it is the signal.

This reframing has a precedent. That the variability in the speech stream is lawful rather than random, conditioned by context, rate, and phonological structure, was recognised decades ago^11^. What has been missing is a neural mechanism that treats that variability predictively, forming an expectation and encoding the deviation from it. A deviation is only informative against a prior: without an expected duration there is no surprisal, and without surprisal there is nothing to read. Prediction and information are thus two sides of one operation: the prior makes the departure computable, and the departure is what carries the information. At the duration level this is literal. The regressor that captured cortical activity is surprisal-based, a quantity defined in the information-theoretic sense, and the cortex was sensitive to it.

Why this matters is easiest to see in cases where timing carries meaning that the identity of the sounds does not: a syllable lengthened to mark emphasis or a phrase boundary, a brief hesitation that signals uncertainty, a fractionally delayed response that reads as reluctance. In each, the information is carried not by what is said but by when, and by how far the timing departs from expectation. Our study does not test the perception of these specific cues; it establishes the mechanism they would require: a cortex that holds temporal expectations and is measurably sensitive to their violation.

One alternative reading should be set aside. That listeners judge duration relatively rather than absolutely is well established: the rate of surrounding speech shifts phonetic category boundaries and can determine whether a word is heard at all^22,23^. Rate normalisation, however, is not prediction. It rescales a decision after the interval has elapsed, drawing on context that may be subsequent to the event being judged; it forms no expectation in advance, and it leaves no quantity behind once the judgement is made. Our regressors are prospective by construction, each computed from intervals that precede the event it scores, and the departure they encode is not consumed by a categorisation but persists as a graded value that predicts cortical activity in its own right.

### Relation to oscillatory accounts: sampling versus coding

The dominant framework for cortical speech processing holds that neural oscillations entrain to the quasi-rhythmic speech envelope, aligning windows of heightened excitability to the syllabic beat so that finer acoustic analysis is sampled where information is dense. Our account should not be read as a rival to this one. The two describe different operations. The literature’s vocabulary tends to obscure the distinction, since “tracking” is applied indifferently to a measurement and to the mechanism presumed to produce it^24^. Entrainment is a sampling mechanism: it phase-aligns processing to a near-periodic carrier, and its temporal expectation is implicit in the frequency of the oscillation itself. It anticipates what recurs. What we describe is an encoding mechanism: it represents the timing of speech events as a quantity, forms an explicit expectation of that timing, and reads the deviation from it as information. That includes events which do not recur periodically at all.

Seen this way, predictive timing does not sit alongside the oscillatory account so much as contains it: entrainment is what adaptive prediction looks like at the one grain where speech approximates periodicity. Where speech timing is locally quasi-regular, on the slow syllabic scaffold, a fixed oscillator suffices. Adaptive prediction at its coarsest grain reduces to something closely resembling entrainment. Nor are oscillatory mechanisms wholly rigid. Theta– gamma coupling adjusts to changes in speech rate^25^, dynamic attending theories implement event-by-event correction of an oscillator’s period and phase^26^, and the segmentation tempo of an oscillatory model can be tuned by contextual predictability^27^. The nearest such account couples an oscillator to linguistic prediction, using the predicted identity of an upcoming word to realign phase where the acoustic signal drifts from periodicity^28^. In each case, however, the departure from expected timing is what the model consumes: a correction signal, absorbed so that the oscillator stays locked, leaving nothing behind for downstream processing to read. Our regressors invert that role. The departure is not cancelled but retained — scored at every event, graded, signed, and read out as a quantity that structures the cortical response. The interval it is computed from is retained alongside it. And the concession may in any case be generous. When syllabic periodicity is allowed to vary naturally rather than being imposed, comprehension improves as periodicity falls, and coupling between posterior superior temporal and speech motor cortex strengthens rather than weakens^7^. That is the opposite of what a mechanism profiting from a stable carrier would predict. Even at the grain where entrainment should be most at home, periodicity as such does not appear to be what listeners exploit. Most of the temporal structure of natural speech lies below that scaffold, in the continuously varying, non-rhythmic timing of individual segments, where an oscillatory mechanism has nothing to lock onto by construction. It is this regime — the aperiodic remainder that periodicity cannot reach — that carries the fine-grained duration and onset information our results show the cortex to encode. Oscillatory sampling and predictive timing are therefore not competing explanations of the same phenomenon but accounts of adjacent ones: the one securing the periodic frame, the other reading the departures from it that periodicity discards.

### Generalisation and scope

The operation we describe — form a local expectation, encode the deviation as information — is not specific to speech. It echoes findings in other domains where the brain is sensitive to violations of temporal expectation: in music, where expressive departures from the metric grid are a principal carrier of structure and performer identity^29^, and where the timing and predictability of events shape perception and neural response^30^, and in vision, where temporal expectation structures attention and processing^31,32^. What these share with the present results is the logic rather than the substrate: a predicted time, an observed departure, and a response scaled to the departure. This suggests that adaptive predictive timing may be a general strategy the brain applies wherever events unfold in time, and that the features it operates over need not be durations. Onsets, intervals, and rates of change are all candidates.

Here we have deliberately narrowed that scope to a single, tractable case: the timing of phonemic events in natural speech, where the departures from expectation are fine-grained, continuous, and consequential for recognition. We make no claim that the same mechanism, in the same form, underlies temporal prediction across domains. What we show is that for speech, the timing of events is not discarded but encoded, predictively, as part of the information the signal conveys. Speech is the domain where the argument is hardest, because its timing is so irregular that it has long been treated as noise.

### Conclusion

Speech-timing research has long treated the temporal irregularity of speech as an obstacle: variance to be normalised, jitter to be tracked through, noise around a rate the brain must recover. Our results invert that premise. Speech unfolds in time not as a carrier to be sampled, but as a structured record of deviations from expectation. The moment-to-moment departure of each event from what was predicted is not what the listening brain discards. It is what the listening brain reads.

## Methods

### Behavioural experiment

Twenty-seven Spanish native speakers completed the behavioural experiment (18 female; mean age = 24.26 years, median = 23 years, range = 19–34 years). Participants were recruited through the Basque Center on Cognition, Brain and Language (BCBL) Participa recruitment system. All were right-handed, reported normal hearing and normal or corrected-to-normal vision, and had no history of language-related impairments. The study was approved by the BCBL Ethics Committee (approval number 031121SM), and all participants provided written informed consent.

Participants heard sentences from the Sharvard Corpus^33^, a phonemically balanced Spanish adaptation of the Harvard sentences, each containing five content words. Of the 700 sentences in the corpus, 693 were used, 231 per condition: natural, isochronous, and anisochronous. Syllabic timing was manipulated following ref. **5**. Syllable onsets were identified using EasyAlign^34^. In the isochronous condition, every inter-syllable interval was set to the sentence mean, producing a regular rhythm; in the anisochronous condition, the same total temporal distortion was randomly redistributed across syllable positions, producing an irregular, non-periodic rhythm. Time-scaling was applied interval by interval using WSOLA^35^, while preserving segmental spectra and F0. Sentences were presented in white noise at −1 dB SNR, a level selected after pilot testing at 0, −1, and −2 dB, and were ramped on and off using raised-cosine ramps of 250 ms and 150 ms, respectively. Stimuli were presented in mini-blocks of three or four sentences from the same condition and pseudo-randomised such that no two consecutive blocks contained the same condition. After each mini-block, participants repeated the final sentence aloud. Comprehension was scored as the proportion of content words correctly recalled, ranging from 0 to 1 in increments of 0.2.

Trials on which the participant produced no response (157 of 5,346, 2.9%; 9 natural, 48 isochronous, 100 anisochronous) were scored as zero keywords recalled rather than excluded, since a failure to report any keyword is an outcome of the task rather than a missing observation, and its frequency varied systematically with condition. Keyword-recall percentages were averaged per participant and condition (66 trials per cell) and entered into a one-way repeated-measures ANOVA with timing condition (natural, isochronous, anisochronous) as a within-participant factor. Sphericity was assessed with Mauchly’s test; as it was not violated, uncorrected degrees of freedom are reported (Greenhouse–Geisser ε = 0.88; the correction did not alter any conclusion). Effect size is reported as generalised eta-squared. The three pairwise comparisons were tested with paired t-tests and Bonferroni-corrected across the family; effect sizes are Cohen’s d_z. Analyses were performed in Python 3 using NumPy and SciPy; the analysis script is available in the repository listed under Code availability.

### MEG experiment

Twenty-five native speakers of Spanish completed the MEG experiment (17 female; mean age = 38.60 years, median = 37.6 years, range = 19–57 years). All participants were right-handed, reported normal hearing and normal or corrected-to-normal vision, and had no history of language-related impairments. The study was approved by the BCBL Ethics Committee (approval number 041121MK), and all participants provided written informed consent in accordance with the Declaration of Helsinki.

### Stimuli

The stimuli consisted of five spontaneous narratives produced by the same female native speaker of Spanish. For each recording, the speaker was given a general topic and asked to speak freely for several minutes without following a script. The five topics were holidays, animals, Switzerland, travel, and wedding. The narratives had a mean duration of 7 min 7 s (SD = 4.7 s). Recordings were made in a sound-attenuated booth at the Basque Center on Cognition, Brain and Language using a Marantz PMD670 digital recorder and digitized at a sampling rate of 44.1 kHz. The resulting WAV files underwent standard noise-reduction procedures and were segmented and annotated at the word and phone levels using Praat.

Word-and phone-level boundaries were generated in TextGrid format automatically using the Montreal Forced Aligner (MFA version 3.0.5)^36^ with the pretrained *spanish_mfa* acoustic model (v2_0_0a) and *spanish_mfa* dictionary (v2_0_0a). The resulting alignments were subsequently inspected and manually corrected by a native Spanish speaker with expertise in phonetics and speech–text alignment.

### Procedure

Participants sat comfortably inside the MEG system in a magnetically shielded room. Throughout auditory presentation, they were instructed to remain still and maintain fixation on a centrally presented fixation point. Before the experiment, the presentation volume delivered through eartubes was adjusted individually to a comfortable and clearly audible level and was subsequently kept constant throughout the session.

Each participant listened to three of the five narratives, selected according to a pseudorandomised assignment procedure. This resulted in approximately 21 min of continuous-speech listening per participant (estimated mean: 21 min 20 s). At the end of each narrative, participants answered six yes/no comprehension questions using the corresponding buttons on an MEG-compatible response device. Thus, each participant answered a total of 18 comprehension questions. Due to a technical error, responses from three participants were not recorded. In addition, a 3-min empty-room recording was acquired. The whole experimental session lasted around one hour, including participant preparation.

### MEG data acquisition and preprocessing

Neuromagnetic activity was recorded using one of two whole-head 306-sensor MEG systems, each comprising 102 magnetometers and 204 planar gradiometers. Fourteen participants were recorded using an Elekta Neuromag Vectorview system and eleven using the subsequently installed MEGIN TRIUX Neo system. The same experimental protocol and nominal acquisition parameters were used for both systems. Signals were recorded with an online passband of 0.1– 330 Hz and sampled at 1 kHz.

Head position within the MEG helmet was continuously monitored using five head-position indicator coils attached to the participant’s scalp. Before recording, the positions of these coils, three anatomical fiducial points—the nasion and left and right preauricular points—and at least 150 additional points distributed over the scalp and nose were digitised in a common coordinate system using a Fastrak electromagnetic tracker (Polhemus).

MEG data were preprocessed using MNE-Python and organised according to the Brain Imaging Data Structure using MNE-BIDS. For recordings acquired with the newer MEG TRIUX system, the nominal sampling frequency was corrected from 1000 to 1000.49 Hz to account for the system-specific clock delay. Flat and noisy channels were first identified automatically using find_bad_channels_maxwell, based on 30-s data windows and requiring detection in at least 10 windows, and were subsequently verified by visual inspection; additional channels were marked manually when necessary. Continuous head-position information was estimated from the head-position indicator coils using 0.5-s windows, and the resulting estimates were used for movement compensation. Maxwell filtering and temporal signal-space separation were then applied using the calibration and cross-talk correction files specific to each MEG system, with a 10-s temporal window and a correlation threshold of 0.98. The resulting data were resampled to 200 Hz, including the corresponding anti-aliasing low-pass filter at 100 Hz.

Physiological artifacts were removed using independent component analysis. A copy of the Maxwell-filtered data was band-pass filtered between 0.5 and 30 Hz and decomposed into 20 independent components using the Picard algorithm. Components associated with horizontal and vertical eye movements, eye blinks, and cardiac activity were identified automatically from their correlations with the horizontal electrooculogram (HEOG), vertical (VEOG), and electrocardiogram (ECG) channels. Component time courses, spatial topographies, correlation scores, and explained variance were visually inspected, and the selected components were adjusted manually when necessary. The identified artifact-related components were then removed from the Maxwell-filtered data before the ICA-specific band-pass filter was applied, thereby preserving the broader frequency content of the recordings for subsequent analyses.

For each participant, the empty-room recording was selected. Any residual digitisation information was removed, and flat and noisy sensors were identified using the same automatic and manual procedures applied to the participant recordings. Empty-room data were processed with machine-specific calibration and cross-talk files using Maxwell filtering and temporal signal-space separation in MEG-device coordinates, with a 10-s window and a correlation threshold of 0.98, and were subsequently resampled to 200 Hz.

High-resolution structural T1-weighted images were acquired using the 3-T Siemens MAGNETOM Prismafit MRI scanner available at the BCBL. Individual cortical brain surfaces were reconstructed from each participant’s high-resolution T1-weighted MRI using the recon-all pipeline implemented in FreeSurfer version 7.4. A three-layer boundary-element model was generated from the anatomical reconstruction using the watershed procedure in MNE-Python, and dense scalp surfaces were created to support MEG–MRI alignment. The digitised anatomical fiducials, head-position indicator coils, and additional scalp and nose points were then manually coregistered to the participant’s anatomical MRI using the MNE-Python coregistration interface. Initial alignment was based on the nasion and bilateral preauricular points and was subsequently refined by matching the digitised head-shape points to the reconstructed scalp surface. A head-to-MRI transformation was saved for each participant.

### Predictor Variables

#### Acoustic baseline

The acoustic representation followed ref. **13**. A gammatone spectrogram was computed over 8 frequency bands with centre frequencies log-spaced from 165 Hz to 11.7 kHz, sampled at 1000 Hz. An accompanying acoustic-onset representation was derived from the same 8-band auditory spectrogram using the half-wave-rectified onset detector of ref. **13**, yielding two 8-band predictors (spectrogram and onsets) entered together as the acoustic control. A phoneme-onset regressor — a unit impulse at each phoneme onset, carrying no magnitude information — was included alongside them to capture the evoked response to phoneme segment onsets independently of any acoustic or linguistic weighting.

#### Linguistic surprisal baseline

Three phoneme-level linguistic surprisal predictors were computed following ref. **13**, indexing the predictability of upcoming speech at increasing contextual scope: sublexical, word, and sentence. Forced-aligned phoneme sequences were mapped to the pronunciation-dictionary inventory and stress markers were removed before analysis. Sublexical surprisal was the negative base-2 logarithm (in bits) of each phoneme’s conditional probability under a 5-gram phoneme model, given the four preceding phonemes across word boundaries; the first phoneme of each narrative segment, having no preceding context, was assigned zero. Word-level surprisal was derived from a cohort model over a Spanish pronunciation lexicon weighted by unigram word frequencies: as each phoneme of a word arrives, the model maintains the set of lexical candidates consistent with the input so far, and surprisal quantifies the reduction in probability mass that the incoming phoneme imposes on the surviving cohort. Sentence-level surprisal used the same cohort computation but with each word’s lexical prior re-weighted by its probability under a 5-gram word model given the preceding context, estimated with KenLM^37^ from the same corpus; contextually predictable words therefore entered the cohort with higher prior probability and incurred lower surprisal. The 5-gram phoneme and word models were estimated with KenLM from the Spanish OpenSubtitles corpus^38^. All three predictors were entered as impulse regressors at phoneme onsets, weighted by their surprisal value in bits; the cohort-and phoneme-entropy measures carried in the same pickle files were not used in the present study.

### Timing predictors

The three timing predictors were derived from Montreal Forced Aligner phoneme and word alignments and share a general expectation-and-comparison logic, although their numerical forms differ. Levels 1 and 2 quantify onset timing as signed, locally adaptive innovations: the difference in milliseconds between the observed event time and an expectation formed from recent intervals, negative when an event arrives earlier than expected and positive when later. Both were entered as impulses at phoneme onsets. Level 3 quantifies the unexpectedness of a completed phoneme duration under an adaptively updated, class-and context-conditioned log-normal model and was represented at phoneme offsets as a negative log-density in nats.

*Levels 1 and 2 — syllabic rhythm adaptation and phoneme onset prediction.* Let o1, o_2_, …, o_n_ denote the ordered event onsets—vowel onsets for Level 1, as a proxy for syllabic nuclei, and all phoneme onsets for Level 2—and let I_κ_= o_κ+1_ − o_k_ denote the corresponding inter-onset intervals. Local pace was tracked using an exponential moving average, with smoothing constant α = 0.2, corresponding to a characteristic memory of approximately five intervals.

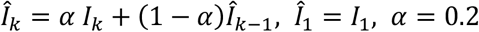

The value assigned to event (κ+1) was the difference between the observed interval and its updated exponential estimate:

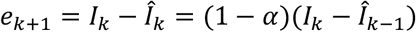

The implemented value is therefore a scaled innovation proportional to the strictly prospective error relative to the preceding interval estimate: early arrivals yield negative values and late arrivals positive values. Values were expressed in milliseconds and clipped at the 95th percentile of their absolute value within each narrative segment to limit the influence of outliers.

The two levels differ only in the event sequence to which the calculation was applied. Level 2 assigns each phoneme its own onset innovation over the complete phoneme-onset sequence. For Level 1, the calculation was applied to vowel onsets, and each resulting value was carried forward across the intervening phonemes until the next vowel. The Level-1 predictor therefore changed only at vowel onsets while providing a value at every phoneme onset. The first event of each sequence was assigned zero because it had no preceding interval; the second was also zero because the first observed interval initialised the moving average.

To test whether the cortical response depends on the direction of the departure and not only its size, the Level-2 innovation was decomposed into two non-negative regressors, both entered as impulses at phoneme onsets: an early-arrival regressor, equal to the magnitude of the departure when the phoneme arrived earlier than expected and zero otherwise, and a late-arrival regressor, defined symmetrically for later-than-expected arrivals. Each event activates exactly one of the two. Their difference recovers the signed innovation and their sum its absolute value, so the pair spans the same information as the signed regressor while allowing the two directions to take separate response functions. The unsigned form used for comparison was the absolute value of the same departure. Across the five narratives, 70% of departures were early and 30% late. The distribution is right-skewed, so symmetric clipping at the 95th percentile of the absolute value truncates more late than early arrivals, leaving the late-arrival regressor the more compressed of the two; this works against, rather than for, detecting a distinct late response. Signed and absolute values nonetheless correlated only weakly (pooled r = 0.11), so direction and magnitude are separable at this grain. Level 1 was not decomposed in this way: there the two correlate at r = 0.49, and direction and magnitude are not separable.

A raw inter-onset interval regressor was also constructed, carrying the interval between each phoneme onset and the preceding one, entered as an impulse at the same event times as the Level-2 departure. The first event of each recording was assigned zero. Because the two regressors are placed at identical event times, they can be compared with timing held constant, as for raw duration and duration surprisal at Level 3. The two are substantially correlated (r = 0.68 across the five narratives), so each nested comparison asks whether one explains cortical activity that the other, despite that shared variance, does not.

*Level 3 — phoneme duration prediction.* Each non-silent phoneme’s duration dᵢ (offset minus onset, in milliseconds) was evaluated against a category-and context-conditioned expectation learned incrementally across the narrative. Phonemes were assigned to a broad class (vowel, stop, fricative, affricate, nasal, liquid or glide), with unmapped labels assigned to an Other fallback class, and further conditioned on an acoustically inferred word-internal stress proxy and phrase-final position, defining a cell c(i). In words containing a single vowel, that vowel was marked as stressed; in words containing multiple vowels, stress was assigned to the vowel maximising the summed z-scores of duration, intensity and pitch. Phrase-final position was inferred from a following silent interval of at least 150 ms, with the final word of the recording also treated as phrase-final. Within each cell, log-durations were modelled as Gaussian with parameters updated online from a common weak prior (μ₀ = ln 80, σ₀ = 0.8, prior weight *κ*₀ = 3 pseudo-observations). Each cell was updated independently, without backoff to broader class models. After n durations had been observed in cell c, with m the mean and v the variance of their logs, the plug-in parameters were

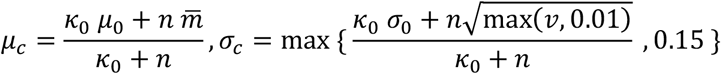

For n ≥ 2, the parameters were updated as specified above. For n = 0, the prior parameters were returned unchanged; for n = 1, σ_c_ = σ_0_ κ_0_/(κ_0_ + 1). The Level-3 value was the negative natural logarithm of the log-normal probability density of the observed duration under the current cell parameters:

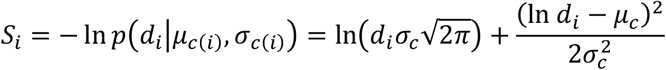

Values were expressed in nats. Because the linguistic-surprisal predictors were expressed in bits, the two scales differ by a constant factor of ln 2; this scaling is absorbed by the TRF coefficients and does not affect model comparison. Crucially, Sᵢ was computed using the model state before dᵢ was observed, and the cell statistics were updated only afterwards, so no phoneme contributed to its own expected-duration model. The predictor was placed at the phoneme offset, when its completed duration first became available. The estimator was reinitialised for each narrative TextGrid, and silent intervals were excluded.

We note two properties of this estimator to avoid over-interpretation. First, it is an online shrinkage-based plug-in estimator, not a full Bayesian posterior-predictive model or a maximum-a-posteriori estimator. The Level-3 value is therefore the negative log-density of the observed duration under the current parameter estimates, rather than an integral over parameter uncertainty. Second, because phoneme duration contributes to the acoustic stress score used to define the class–context cell, stress conditioning is not fully independent of the modelled quantity. In a separate robustness analysis, recomputing Level 3 with stress inferred from intensity and pitch alone left the predictor essentially unchanged (per-segment Pearson r = 0.98–0.99 across the five segments; n = 24,138 phonemes), indicating that this dependence was negligible in practice.

### Multivariate temporal response function analyses

Source-localised MEG responses were analysed using multivariate temporal response functions (mTRFs), following the general framework described by ref. **13**. Analyses were implemented using Eelbrain 0.41.2 and TRF-Tools 11 and were run on the Hyperion high-performance computing cluster at the Donostia International Physics Center. Cross-hemispheric comparisons additionally used a modified build, described below. In an mTRF model, each continuous or discrete stimulus predictor is convolved with a corresponding temporal response function, and the resulting partial responses are summed to predict the continuous neural response. All predictors were fitted jointly, allowing the response associated with each predictor to be estimated while accounting for variance shared with the remaining predictors.

MEG data and continuous stimulus predictors were band-pass filtered between 1 and 20 Hz. Source activity was estimated using a fixed-orientation minimum-norm estimate with an assumed signal-to-noise ratio of 6, corresponding to a regularisation parameter of λ² = 1/36, without depth weighting. Noise covariance matrices were estimated from the empty-room recordings. The cortical source space was defined using an ico-4 tessellation, yielding 2,562 vertices per hemisphere, and individual source estimates were morphed to the FreeSurfer fsaverage surface for group-level analysis. Analyses were restricted to bilateral auditory temporal cortex, defined anatomically as the transverse temporal and superior temporal regions of the Desikan–Killiany parcellation^39^.

Neural responses and stimulus predictors were resampled to 100 Hz, and TRFs were estimated over time lags ranging from −300 to 800 ms. Models were fitted using a boosting algorithm that iteratively minimised the squared prediction error. The continuous recordings were divided into eight contiguous temporal partitions for cross-validation, and selective stopping was used to regularise model fitting and limit overfitting. The same partitions and fitting parameters were used for all models entering a given comparison.

The baseline model included an eight-band gammatone spectrogram, an eight-band acoustic-onset spectrogram, and a phoneme-onset predictor. Linguistic processing was controlled for using phoneme-level surprisal estimates derived from sublexical, word-level, and sentence-level contexts. The predictors of primary interest characterised temporal prediction at three levels: syllabic-rhythm prediction, phoneme-onset timing prediction, and phoneme-duration prediction. The unique contribution of each timing predictor was evaluated by comparing a full model containing that predictor with an otherwise identical reduced model from which the predictor had been omitted. Because model performance was evaluated on held-out data, positive differences indicated that the predictor explained neural-response variance that could not be accounted for by the acoustic, phoneme-onset, linguistic-surprisal, and remaining timing predictors.

Additional nested comparisons tested whether phoneme-duration surprisal explained neural activity beyond raw phoneme duration and, conversely, whether raw duration explained additional variance after accounting for duration surprisal. Further nested comparisons at Level 2 tested whether the direction of the onset departure contributed beyond its magnitude, whether the late-arrival regressor contributed beyond the early-arrival regressor, and whether the departure and the raw inter-onset interval each explained variance beyond the other. Two additional non-nested comparisons contrasted the signed against the unsigned form of the Level-2 regressor, and the departure against the raw inter-onset interval. All models entering a given comparison carried identical acoustic, phoneme-onset, linguistic-surprisal and remaining timing controls, and used the same partitions and fitting parameters. As above, nested comparisons were tested one-tailed and non-nested comparisons two-tailed. A final non-nested comparison contrasted a model containing the three timing-prediction variables with a model containing the three linguistic-surprisal variables, while retaining identical acoustic and phoneme-onset controls.

### Statistical analysis of model comparisons

Model predictive performance was quantified at each cortical source using det, which indexes the proportion of determinable neural-response variance explained in held-out data. For each participant and nested comparison, the unique contribution of the predictor of interest was quantified as the difference between the full and reduced models (Δdet = det_full − det_reduced). Positive Δdet values indicated that including the predictor improved the prediction of unseen neural data.

Group-level inference was performed using related-samples t-tests comparing the full-and reduced-model det maps across the 25 participants. One-tailed tests were used for the planned nested comparisons, testing the directional hypothesis that the full model explained more determinable variance than the reduced model. Multiple comparisons across vertices within the bilateral superior temporal mask were controlled using threshold-free cluster enhancement (TFCE^40^), with significance assessed using 10,000 within-participant permutations and a TFCE-corrected threshold of p < .05.

The three timing predictors represented separate a priori hypotheses, and correction was applied across cortical sources within each statistical test. Bonferroni-corrected p-values across the three model comparisons are additionally reported in the Results. The non-nested comparison between the timing-prediction and linguistic-surprisal models was evaluated using a two-tailed related-samples test, because either model could show greater predictive performance.

Hemispheric differences were assessed in two ways. Differences in predictive contribution were tested by morphing each participant’s Δdet map to the FreeSurfer *fsaverage_sym* surface and comparing homologous vertices across hemispheres within the superior temporal mask, using two-tailed related-samples t-tests with the same TFCE procedure and 10,000 permutations. Differences in response amplitude were tested on the temporal response functions themselves, averaged across sources within the mask separately for each hemisphere, using two-tailed related-samples t-tests over the −100 to 600 ms lag range with TFCE correction across time and 10,000 permutations. Each level was treated as a separate a priori hypothesis and tested individually. Because source orientations are fixed relative to the cortical normal and the two masks differ in extent and gyral geometry, between-hemisphere amplitude comparisons are interpreted with caution; the Δdet comparison, which is invariant to source sign, provides the more conservative test. Both hemispheric analyses were run in a modified Eelbrain build that supports cross-hemispheric morphing; the modification is not reflected in the reported version string, and the build is archived in the code record.

## Data availability

The data underlying Figures 1–5 and Table 1 are available at https://doi.org/10.5281/zenodo.21922711. This record contains the behavioural trial-level data, and the forced-aligned annotations and derived timing and linguistic predictors for the five narratives. The narrative recordings and their alignments are deposited at https://doi.org/10.5281/zenodo.21922092. The stimulus materials for the behavioural experiment derive from the Sharvard Corpus^33^, available at https://doi.org/10.5281/zenodo.3547446. Raw magnetoencephalography recordings and structural MRI scans are not publicly deposited: they are potentially identifiable, participant consent covered controlled scientific reuse but not open publication, and the data must remain within the European Union. De-identified raw data are available to qualified researchers from the corresponding author under a data-use agreement approved by the BCBL Ethics Committee, and will be transferred through European infrastructure. Requests will be answered within 5 weeks.

## Code availability

All analysis code is archived at https://doi.org/10.5281/zenodo.21922711 and https://doi.org/10.5281/zenodo.21922092. This includes the two-stage pipeline that generates the timing predictors from forced-aligned annotations, the scripts that build the rectified Level 2 regressors and the raw inter-onset interval regressor, the departure diagnostics, the behavioural analysis script, and the scripts and notebook used to fit the temporal response function models and to produce the figures. Analyses used Eelbrain 0.41.2, TRF-Tools version 11, MNE-Python 1.9.0, NumPy 2.2.6, SciPy 1.14.1 and KenLM 0.3.0 under Python 3.11. The cross-hemispheric analyses used a locally modified Eelbrain build derived from 0.41.2, which reports the same version string; that source tree is included in the code record. TRF-Tools was installed from the project repository’s development branch rather than a tagged release; the exact source archive used is identified by sha256 checksum 56af3badf9cfdf7cd143432ab8770e6541938f38673cacb7d3e88d1fa7bad6e4 and a copy of that source tree is included in the Zenodo code record. The complete computational environment is provided there as a Conda environment specification (environment_atp.yml) and a full package listing with build strings (environment_atp_versions.txt).

## Author contributions

N.M. conceived the study and developed the theoretical framework. N.M. and J.P.-N. designed the experiments. J.P.-N. collected and analysed the behavioural data. N.M. and J.P.-N. collected the MEG data. N.M. developed the timing predictors and performed the temporal response function analyses. N.M. and J.P.-N. wrote the manuscript. Both authors reviewed and approved the final version.

## Acknowledgements

We thank the participants for their time, and Amets Esnal for recording the narrative stimuli. We thank José Antonio Gonzalo for manual correction of the forced alignments. We thank the members of the mTRF meetings, the Brain Rhythms and Cognition group and the whole BCBL environment. Computational resources were provided by the BCBL and the Donostia International Physics Center (DIPC).

## References

1. Giraud, A.-L. & Poeppel, D. Cortical oscillations and speech processing: emerging computational principles and operations. Nat. Neurosci. 15, 511–517 (2012).

2. Poeppel, D. & Assaneo, M. F. Speech rhythms and their neural foundations. Nat. Rev. Neurosci. 21, 322–334 (2020).

3. Ding, N. & Simon, J. Z. Cortical entrainment to continuous speech: functional roles and interpretations. Front. Hum. Neurosci. 8, 311 (2014).

4. Varnet, L., Ortiz-Barajas, M. C., Guevara Erra, R., Gervain, J. & Lorenzi, C. A cross-linguistic study of speech modulation spectra. J. Acoust. Soc. Am. 142, 1976–1989 (2017).

5. Aubanel, V. & Schwartz, J.-L. The role of isochrony in speech perception in noise. Sci. Rep. 10, 19580 (2020).

6. Aubanel, V., Davis, C. & Kim, J. Exploring the role of brain oscillations in speech perception in noise: intelligibility of isochronously retimed speech. Front. Hum. Neurosci. 10, 430 (2016).

7. Kwon, S., Lubinus, C., Kell, C. A., Keitel, A. & Rimmele, J. M. Effects of speech periodicity and speech rate on auditory-motor coupling during speech comprehension. Commun. Biol. 9, 205 (2026).

8. Klatt, D. H. Linguistic uses of segmental duration in English: acoustic and perceptual evidence. J. Acoust. Soc. Am. 59, 1208–1221 (1976).

9. Aylett, M. & Turk, A. The smooth signal redundancy hypothesis: a functional explanation for relationships between redundancy, prosodic prominence, and duration in spontaneous speech. Lang. Speech 47, 31–56 (2004).

10. Jaeger, T. F. Redundancy and reduction: speakers manage syntactic information density. Cogn. Psychol. 61, 23–62 (2010).

11. Elman, J. L. & McClelland, J. L. Exploiting lawful variability in the speech wave. In Invariance and Variability in Speech Processes 360–385 (Erlbaum, 1986).

12. Deyna, L. et al. Cortical encoding of probabilistic temporal predictions during speech perception. bioRxiv 10.64898/2026.08.16.745095 (2026).

13. Brodbeck, C. et al. Parallel processing in speech perception with local and global representations of linguistic context. eLife 11, e72056 (2022).

14. Heilbron, M., Armeni, K., Schoffelen, J.-M., Hagoort, P. & de Lange, F. P. A hierarchy of linguistic predictions during natural language comprehension. Proc. Natl Acad. Sci. USA 119, e2201968119 (2022).

15. Nolan, F. & Jeon, H.-S. Speech rhythm: a metaphor? Phil. Trans. R. Soc. B 369, 20130396 (2014).

16. Grabe, E. & Low, E. L. Durational variability in speech and the Rhythm Class Hypothesis. In Laboratory Phonology 7 (eds Gussenhoven, C. & Warner, N.) 515–546 (Mouton de Gruyter, 2002).

17. Low, E. L., Grabe, E. & Nolan, F. Quantitative characterizations of speech rhythm: syllable-timing in Singapore English. Lang. Speech 43, 377–401 (2000).

18. Breska, A. & Ivry, R. B. Double dissociation of single-interval and rhythmic temporal prediction in cerebellar degeneration and Parkinson’s disease. Proc. Natl Acad. Sci. USA 115, 12283–12288 (2018).

19. Buzsáki, G. Time, space, memory and brain–body rhythms. Nat. Rev. Neurosci. (2025). doi:10.1038/s41583-025-00987-2

20. Poeppel, D. The analysis of speech in different temporal integration windows: cerebral lateralization as ‘asymmetric sampling in time’. Speech Commun. 41, 245–255 (2003).

21. Oderbolz, C., Poeppel, D. & Meyer, M. Asymmetric sampling in time: evidence and perspectives. Neurosci. Biobehav. Rev. 171, 106082 (2025).

22. Miller, J. L. & Liberman, A. M. Some effects of later-occurring information on the perception of stop consonant and semivowel. Percept. Psychophys. 25, 457–465 (1979).

23. Dilley, L. C. & Pitt, M. A. Altering context speech rate can cause words to appear or disappear. Psychol. Sci. 21, 1664–1670 (2010).

24. Molinaro, N. The tracking umbrella: diverse interpretations under a common neural term. Ann. N. Y. Acad. Sci. 1552, 36–46 (2025).

25. Lizarazu, M. et al. Phase–amplitude coupling between theta and gamma oscillations adapts to speech rate. Ann. N. Y. Acad. Sci. 1453, 140–152 (2019).

26. Large, E. W. & Jones, M. R. The dynamics of attending: how people track time-varying events. Psychol. Rev. 106, 119–159 (1999).

27. Platonova, O., Dogonasheva, O., Giraud, A.-L. & Bouton, S. Contextual prediction tunes the tempo of speech segmentation. bioRxiv 10.64898/2026.03.31.713600 (2026).

28. ten Oever, S. & Martin, A. E. An oscillating computational model can track pseudo-rhythmic speech by using linguistic predictions. eLife 10, e68066 (2021).

29. Repp, B. H. A microcosm of musical expression. I. Quantitative analysis of pianists’ timing in the initial measures of Chopin’s Etude in E major. J. Acoust. Soc. Am. 104, 1085–1100 (1998).

30. Bianco, R. et al. Human newborns form musical predictions based on rhythmic but not melodic structure. PLOS Biol. (2026). doi:10.1371/journal.pbio.3003600

31. Nobre, A. C. & van Ede, F. Anticipated moments: temporal structure in attention. Nat. Rev. Neurosci. 19, 34–48 (2018).

32. Shalev, N., Nobre, A. C. & van Ede, F. Time for what? Breaking down temporal anticipation. Trends Neurosci. 42, 373–374 (2019).

33. Aubanel, V., García Lecumberri, M. L. & Cooke, M. The Sharvard Corpus: a phonemically-balanced Spanish sentence resource for audiology. Int. J. Audiol. 53, 633–638 (2014).

34. Goldman, J.-P. EasyAlign: an automatic phonetic alignment tool under Praat. In Proc. Interspeech 2011, 3233–3236 (2011).

35. Demol, M., Verhelst, W., Struyve, K. & Verhoeve, P. Efficient non-uniform time-scaling of speech with WSOLA. In Proc. SPECOM 2005, Vol. 1, 163–166 (2005).

36. McAuliffe, M., Socolof, M., Mihuc, S., Wagner, M. & Sonderegger, M. Montreal Forced Aligner: trainable text-speech alignment using Kaldi. In Proc. Interspeech 2017, 498–502 (2017).

37. Heafield, K. KenLM: faster and smaller language model queries. In Proc. Sixth Workshop on Statistical Machine Translation 187–197 (Association for Computational Linguistics, 2011).

38. Lison, P., Tiedemann, J. & Kouylekov, M. OpenSubtitles2018: statistical rescoring of sentence alignments in large, noisy parallel corpora. In Proc. LREC 2018 (European Language Resources Association, 2018).

39. Desikan, R. S. et al. An automated labeling system for subdividing the human cerebral cortex on MRI scans into gyral based regions of interest. NeuroImage 31, 968–980 (2006).

40. Smith, S. M. & Nichols, T. E. Threshold-free cluster enhancement: addressing problems of smoothing, threshold dependence and localisation in cluster inference. NeuroImage 44, 83–98 (2009).

